# Using extreme gradient boosting (XGBoost) to evaluate the importance of a suite of environmental variables and to predict recruitment of young-of-the-year spotted seatrout in Florida

**DOI:** 10.1101/543181

**Authors:** Elizabeth Herdter Smith

## Abstract

Environmental factors strongly influence the success of juvenile fish recruitment and productivity, but species-specific environment-recruitment relationships have eluded researchers for decades. Most likely, this is because the environment-recruitment relationship is nonlinear, there are multi-level interactions between factors, and environmental variability may differentially affect recruitment among populations due to spatial heterogeneity. Identifying the most influential environmental variables may result in more accurate predictions of future recruitment and productivity of managed species. Here, gradient tree boosting was implemented using XGBoost to identify the most important predictors of recruitment for six estuary populations of spotted seatrout (*Cynoscion nebulosus*), an economically valuable marine resource in Florida. XGBoost, a machine learning method for regression and classification, was employed because it inherently models variable interactions and seamlessly deals with multi-collinearity, both of which are common features of ecological datasets. Additionally, XGBoost operates at a speed faster than many other gradient boosting algorithms due to a regularization factor and parallel computing functionality. In this application of XGBoost, the results indicate that the abundance of pre-recruit, juvenile spotted seatrout in spatially distinct estuaries is influenced by nearly the same set of environmental predictors. But perhaps of greater importance is that the results of this study show that this algorithm is highly effective at predicting species abundance and identifying important environmental factors (i.e. predictors of recruitment). It is strongly encouraged that future research explore the applicability of the XGBoost algorithm to other topics in marine and fisheries science and compare its performance to that of other statistical methods.

## Introduction

Environmental factors such as ENSO cycles, salinity, and temperature can strongly influence the success of fish recruitment worldwide (Hjort 1914; Cushing 1982); however the true environment-recruitment relationship is difficult to identify. Several reasons for this difficulty are described by Rose (2000) and Keyl and Wolff (2008) and summarized here. First, the relationship between environmental factors and recruitment is often nonlinear, and many commonly used statistical methods are unable to model such nonlinearity. Additionally, environmental factors may interact to limit or enhance the recruitment response or mitigate or enhance the effects of other environmental factors. Moreover, environmental variability may differentially affect recruitment among populations or subpopulations due to the spatial heterogeneity of the influence of environmental factors. Further, researchers often evaluate the influence of environmental variables at spatial scales that are incongruous with the spatial scale of recruitment. Finally, an environment-recruitment relationship may change due to a shift in the environmental regime (Rose 2000; Keyl and Wolff 2008; Vert-pre et al. 2013).

Nevertheless, identifying environmental variables that are most influential to recruitment and including these sources of variability in the estimation process may yield more accurate estimates of current and future recruitment (Myers 1998). Moreover, accurate estimates of historical and future recruitment are vital to the sustained productivity of exploited populations because parameters that describe the stock-recruitment relationship are used to calculate biological reference points and fishing mortality targets (Hilborn and Walters 1992). Moreover, identifying highly influential environmental variables can inform adaptive management strategies that accommodate for changes in the environment. Thus, this task is of paramount importance (Collie et al. 2016).

In Florida, annual recruitment of spotted seatrout (*Cynoscion nebulosus*), an estuarine fish of great economic importance to the recreational fishery in the Gulf of Mexico, is characterized by deviations unexplainable by changes in adult biomass. In addition, poor fits of Beverton-Holt and Ricker stock-recruitment curves to adult and young-of-the-year (YOY) abundance is suggestive of an unaccounted for source of process error, likely an environmental factor, in the quantitative assessment of this stock (Murphy et al. 2011). Growth rates and recruitment strength differ among genetically similar but spatially separate estuary populations of spotted seatrout, which is indicative of the likely influence of local environmental variables on recruitment strength of this species (Herdter et al. in review, Murphy and McMichael 2003; Kupschus 2004). Further, previous studies have found significant correlations between the abundance of YOY spotted seatrout in several estuaries in Florida and environmental variables, such as freshwater inflow, sea surface temperature, salinity and seagrass cover (Peebles and Tolley 1988; Holt and Holt 2003; Matheson et al. 2003; Kupschus 2004; Purtlebaugh and Allen 2010; Dutterer et al. 2013; Flaherty-Walia et al. 2015).

Such studies are extremely valuable in the quest to identify an environment-recruitment relationship for spotted seatrout in Florida. However, most researchers have employed traditional statistical methods such as generalized linear modeling or correlation analyses, which assume that the environment-recruitment relationship is stationary over time. In contrast, the true relationship may be nonlinear or non-monotonic in nature; thus, the statistical methods used in the aforementioned studies may have resulted in large type I or type II errors (Keyl and Wolff 2008). Additionally, these studies used only short-term data sets and did not exhaustively examine possible temporal lags; thus, the true relationships or temporal lags may have been undetected. To reduce the chance of identifying a spurious environment-recruitment relationship, the underlying mechanism of such relationships must be defined and supported by cross-validation, all possible environmental predictors must be screened, and the final relationship must be tested on independent data (Myers 1998; Francis 2006).

The objective of this study was to build predictive models of recruitment for six estuary populations of spotted seatrout in Florida using extreme gradient boosting (XGBoost), a new gradient boosted regression tree algorithm developed by Chen and Guestrin (2016), to illuminate spatial differences in predictor importance. Gradient boosted regression trees were employed because they inherently model variable interactions, incorporate *k*-fold cross-validation into the training algorithm and evaluate the predictive model using a withheld testing dataset (De’ath and Fabricius 2000; Friedman 2001; Elith et al. 2008). This research expands on the current understanding of an environment-recruitment relationship for spotted seatrout in Florida by simultaneously evaluating all available environmental variables at an appropriate spatial scale and considering their influence at several time lags to account for their effects on both spawning success and YOY survival and recruitment. This research provides the foundation for efforts to evaluate environmental variables within a quantitative framework for spotted seatrout in Florida. Such insight is imperative for effective management of this valuable resource.

## Methods

### Spotted seatrout abundance data

Observations of YOY spotted seatrout abundance were used as a proxy for spotted seatrout recruitment and were obtained by the Fisheries Independent Monitoring (FIM) program at the Florida Fish and Wildlife Conservation Commission’s (FWC) Fish and Wildlife Research Institute (FWRI) in St. Petersburg (FWRI 2016). The FIM program surveys six major estuaries in Florida using a multi-gear, stratified-random sampling (SRS) approach. The FIM program routinely monitors the Jacksonville area of northeast Florida (JX) which includes the St. Marys River, Cumberland Sound, the Nassau River and Sound, and the St. Johns River. Apalachicola Bay (AP), Cedar Key (CK), the northern and southern portions of the Indian River Lagoon (Mosquito Lagoon through the Sebastian Inlet and Vero Beach to the Jupiter Inlet; IR), Tampa Bay (TB), and Charlotte Harbor (CH; Figure 1) are also routinely monitored.

**FIGURE 1.**
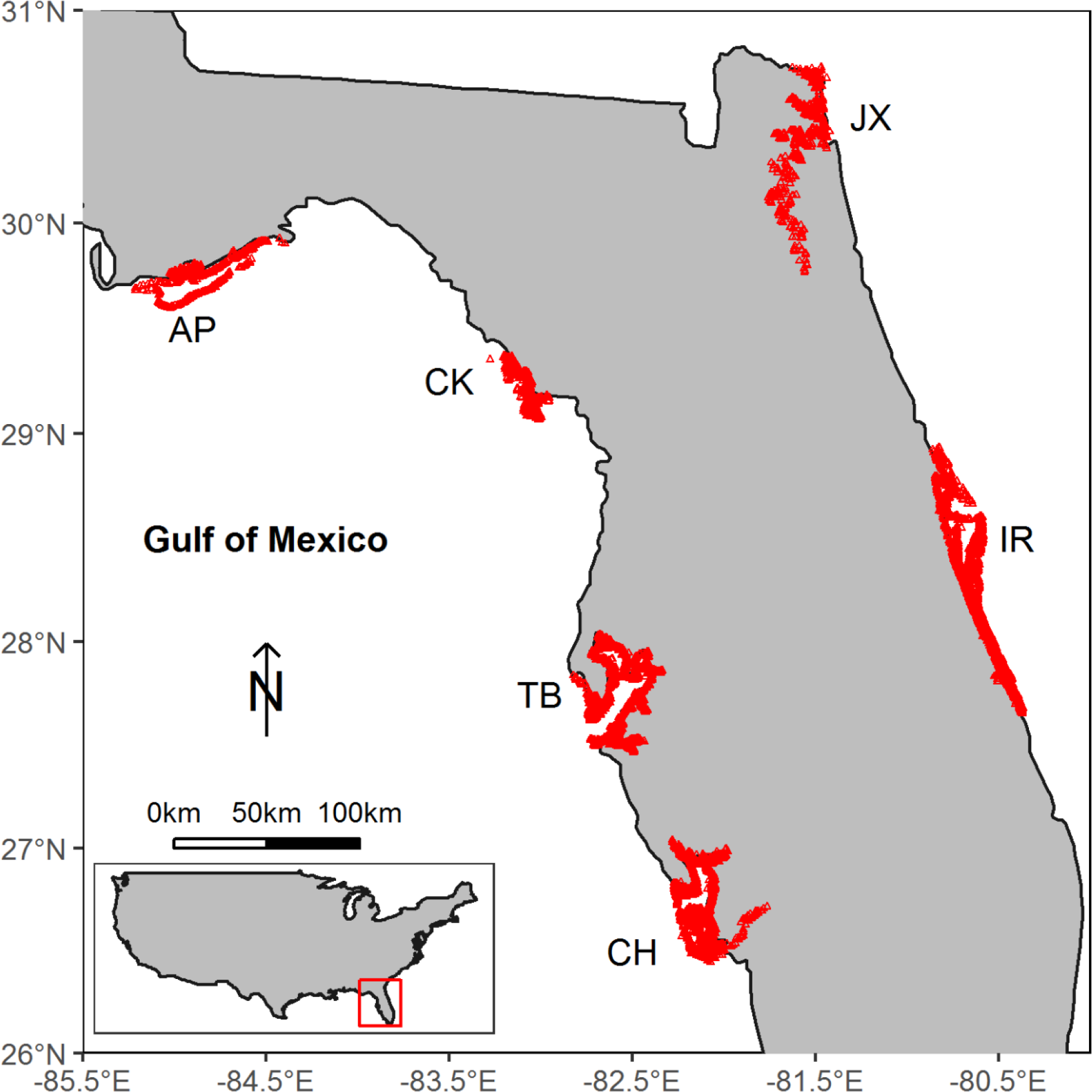
Map of statewide sampling for juvenile spotted seatrout in six sampling estuaries in Florida: Apalachicola (AP), Cedar Key (CK), Tampa Bay (TB), Charlotte Harbor (CH), northeast Florida (JX), and the Indian River Lagoon (IR). The red triangles indicate unique sampling stations across all years within each estuary.

The FIM program started sampling for YOY and juvenile fishes in Tampa Bay and Charlotte Harbor in 1989 using 21.3 - meter bag seines. Monthly sampling efforts expanded to the northern region of the Indian River Lagoon, Cedar Key, Apalachicola Bay, and northeast Florida in 1990, 1996, 1998, and 2001, respectively. In 1996, the sampling program started monitoring larger fish, including juvenile and adult spotted seatrout, with 183-meter haul seines (FWRI, 2016). Observations of YOY abundance from all six estuaries except for data from the southern portion of the Indian River Lagoon were used for this study. The data from the southern Indian River Lagoon were excluded because the FIM program does not sample the southern Indian River Lagoon with a 21.3-meter seine; therefore, there are no estimates of YOY abundance in that area. At every haul, hereafter referred to as a sampling event, the number of YOY spotted seatrout in the seine was recorded (a zero was recorded if none were present), and water quality was assessed with a YSI^®^ water quality meter. Young-of-the-year fish were defined as those less than or equal to 100 millimeters standard length (SL; Murphy et al. 2006).

### Environmental data

Environmental factors hypothesized to affect adult spawning success and survival and subsequent settlement of YOY spotted seatrout in each area were included in the analysis, when available (Table 1). Factors likely to affect adult spawning success include salinity (at-spawn salinity; Brown-Peterson and Thomas 1988; Saucier and Baltz 1993), water temperature (at-spawn water temperature; Lowerre-Barbieri et al. 1999; Brown-Peterson 2003; Kupschus 2004), and water quality (dissolved inorganic nitrogen as a proxy, [DIN 1prior]) in the vicinity of spawning and during months of supposed spawning (Wootton 1998; Lambert and Dutil 2000; Greening and Janicki 2006). However, these variables were included in the analysis for only Tampa Bay, Charlotte Harbor, and the Indian River Lagoon because the spawning locations in these areas have been previously identified (Gilmore 2003; Lowerre-Barbieri et al. 2009; Walters et al. 2009; Table 1). Environmental factors hypothesized to affect survival and settlement of YOY fish include salinity (Holt and Holt 2003; Wuenschel et al. 2004), water temperature (Alsuth and Gilmore 1994; Wuenschel et al. 2004), dissolved oxygen (Dissolved O_2_; Siefert and Spoor 1974; Breitburg et al. 1994; Miller and Kendall 2009), water clarity (attenuation coefficient as a proxy, [Atten coef]; Fiksen et al. 2002), water depth (Bortone 2003), monthly flow rate of the closest river to each sampling event (river flow-CLOSEST), average monthly flow rate of all rivers that drain into the sampling area (river flow ALL; Dutterer et al. 2013), precipitation, the Palmer Z drought index which is an indicator for short-term drought (Z anomaly; King et al. 2003; Dolbeth et al. 2010), and maximum and minimum air temperature anomalies (maximum/minimum T anomaly). In addition, the concentration of dissolved inorganic nitrogen (DIN) was included in the analysis for Tampa Bay, Charlotte Harbor, and the Indian River Lagoon, and chlorophyll *a* (Chlor *a*) concentration was included in the analysis for Tampa Bay. Both DIN and Chlor *a* concentrations serve as proxies for water quality and productivity of trophic levels upon which YOY Spotted seatrout rely but were not available for the remaining estuaries (Livingston 2001; Sigua and Tweedale 2003; Greening and Janicki 2006; Turner et al. 2006). The influence of environmental variability on YOY spotted seatrout recruitment and abundance was analyzed on an estuary-specific level rather than statewide to examine spatial differences in an environment-recruitment relationship.

**TABLE 1.**
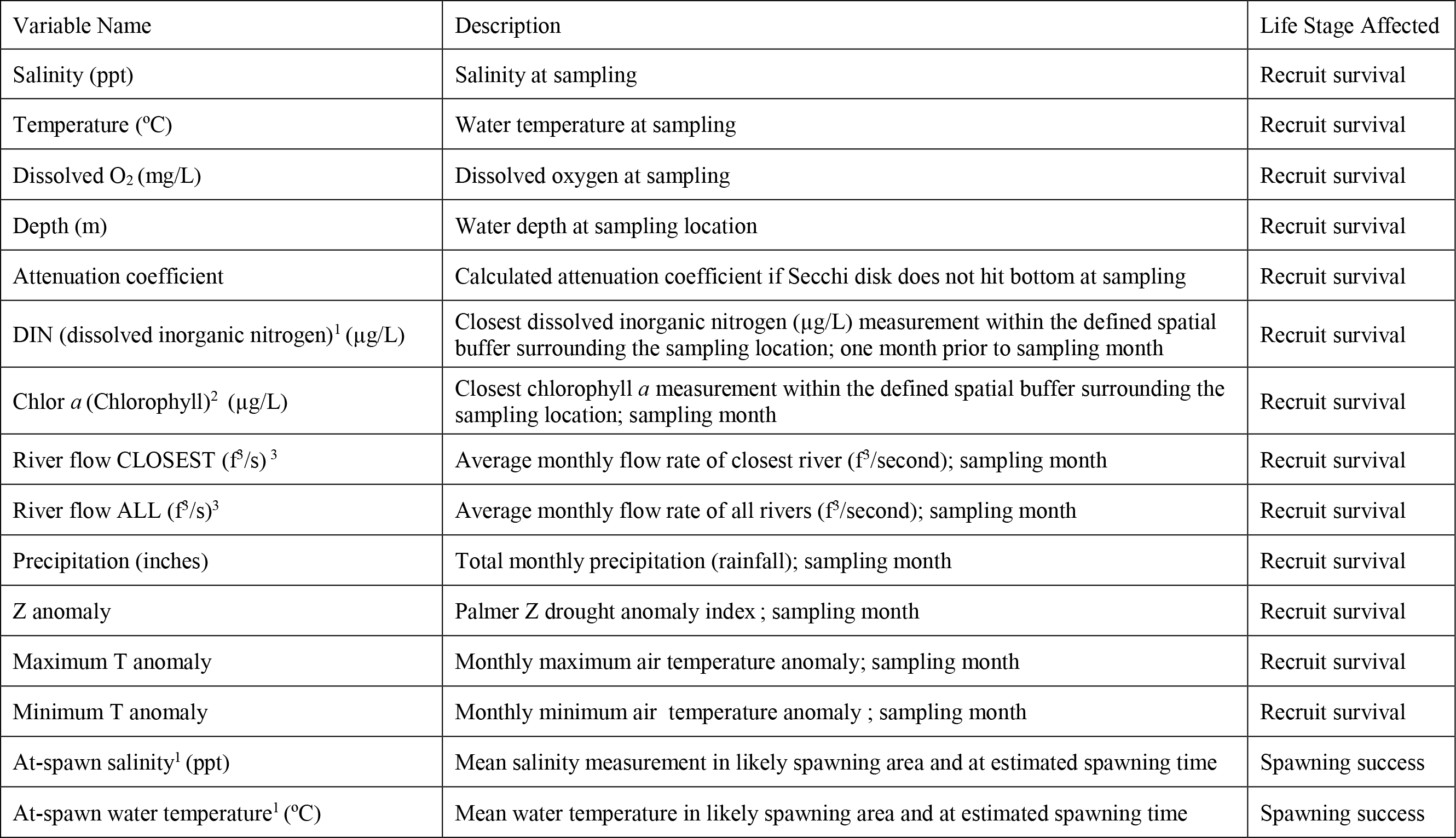
Environmental predictor variables assessed by the XGBoost algorithm and GLM. ^1^Available for Tampa Bay, Charlotte Harbor, and Indian River Lagoon only. ^2^Available for Tampa Bay only. ^3^Apalachicola area has only one major river therefore river flow CLOSEST is equivalent to river flow ALL

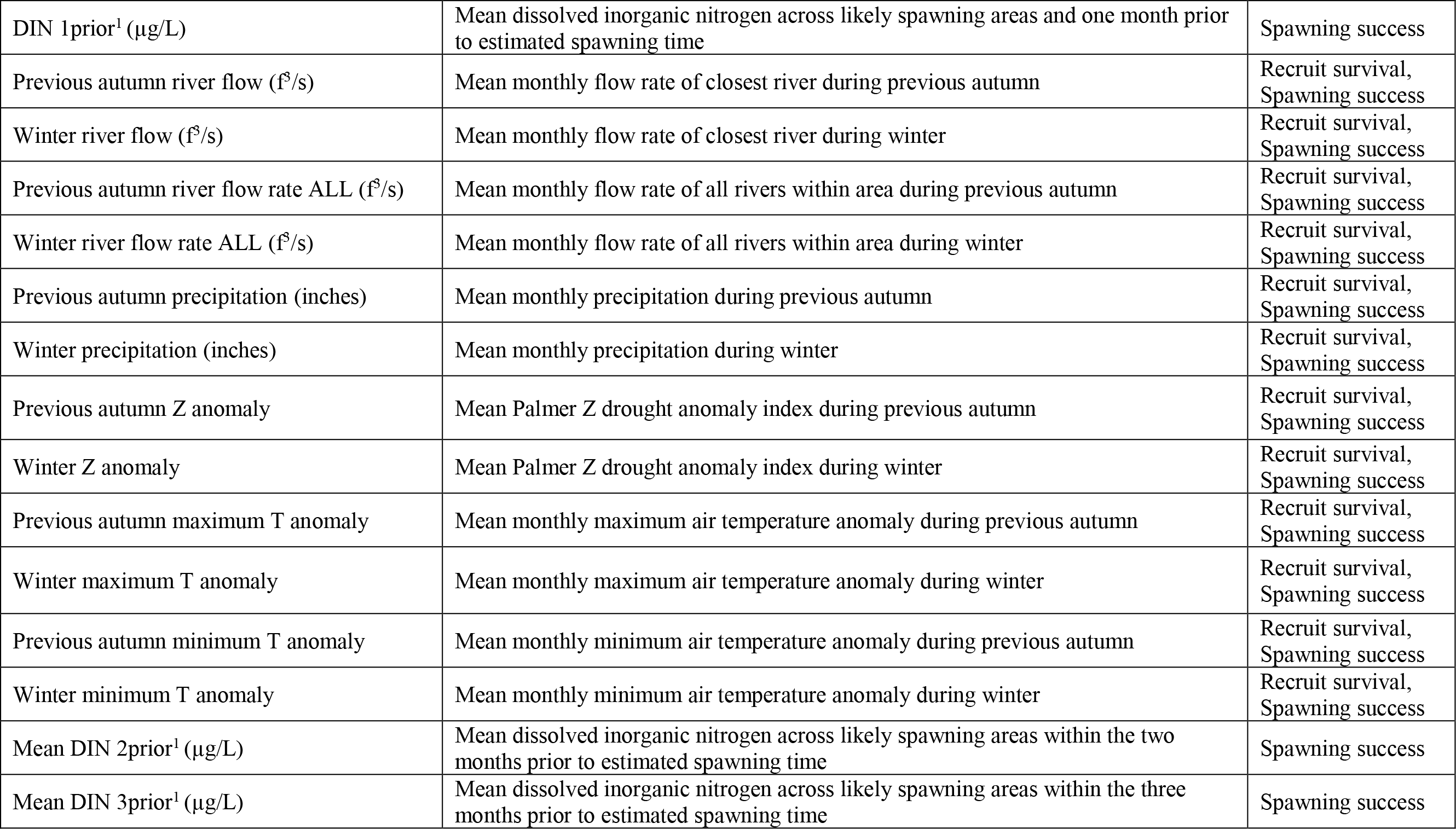

Salinity, water temperature, dissolved oxygen, turbidity, and water depth observations were recorded concomitantly with every sampling event by the FIM program. Nitrogen and chlorophyll *a* concentrations within Tampa Bay were recorded by gauges managed by the Hillsborough County Environmental Protection Commission. Nitrogen concentrations in Charlotte Harbor were recorded by gauges managed by the Lee County Environmental Laboratory, Charlotte Harbor Estuaries Volunteer Water Quality Monitoring Network, and the Peace River Manasota Regional Water Supply Authority. The nitrogen and chlorophyll *a* data were accessed through the Tampa Bay Water Atlas program (http://www.tampabay.wateratlas.usf.edu/datadownload/SelectStations.aspx). Nitrogen concentrations in the Indian River Lagoon were recorded by gauges managed by the St. Johns River Water Management District and were accessed using the data portal on their website (https://www.sjrwmd.com/data/water-qual). The flow rates recorded at the most downstream gauges with the longest data records for each major river within each estuary were obtained from the United States Geological Survey National Water Information System (USGS NWIS; https://maps.waterdata.usgs.gov/mapper/; Table 2). When available, tidally filtered flow rates were preferred over general flow rates if the river was tidally influenced. The Z anomaly, and maximum and minimum T anomalies were obtained for each national climatological division (CD) within which each estuary is located from the National Oceanic and Atmospheric Administration National Centers for Environmental Information (NOAA NCEI; https://www.ncdc.noaa.gov/cag/divisional/ti; Table 2). Precipitation data recorded at the weather station with the longest continuous data record and closest to each estuary were also obtained from NOAA NCEI (https://gis.ncdc.noaa.gov/maps/ncei/s; Table 2).

**TABLE 2.** River gauge numbers, weather station locations, and climate division assignments for each estuary: Apalachicola (AP), Cedar Key (CK), Tampa Bay (TB), Charlotte Harbor (CH), and northeast Florida (JX), and the Indian River Lagoon (IR).

| Area | Rivers | Weather Station | Climate Division |
| --- | --- | --- | --- |
| AP | Apalachicola River (gauge #02359170) | Apalachicola Airport Station: USW00012832 | 1 |
| CK | Suwannee River (station # 02323500)<br>Waccasassa River (station #02313700) | Usher Tower, FL Station: USC00089120 | 2 |
| TB | Alafia River (gauge #2301500)<br>Hillsborough River (gauge #2304500)<br>Little Manatee River (gauge #2300500) | Tampa International Airport, Station:<br>USW00012842 | 4 |
| CH | Caloosahatchee River (gauge #2292900)<br>Myakka River (gauge #2298830)<br>Peace River (gauge #2296750) | Fort Myers Page Field Airport,<br>Station:USW00012835 | 5 |
| JX | Cedar River (gauge #02246459)<br>Ortega River (gauge #02246318)<br>St. Johns River at Buffalo Bluff (gauge #02244040)<br>St. Johns River (gauge #02246500)<br>St. Marys River (gauge #02231254) | Jacksonville Naval Air Station, Station:<br>USW00093837 | 2 |
| IR | Crane Creek (gauge # 02249500)<br>Eau Gallie River (gauge #02249007)<br>North Prong St. Sebastian River (gauge #02251500)<br>South Prong St. Sebastian River (gauge #02251000)<br>Turkey Creek (gauge # 02250030) | Melbourne Weather Forecast Office, Station:<br>USC00085612 | 3 |

The attenuation coefficient, DIN (including DIN 1prior), monthly river flow, and monthly precipitation were derived before they were included as environmental variables in the estuary-specific datasets. The attenuation coefficient was not available in the FIM dataset so it was calculated by dividing the Secchi depth recorded concomitantly with every sampling event by 1.7 to obtain the attenuation coefficient, *k* (Murty 1969). However, the attenuation coefficient was not calculated if the Secchi disk reached the bottom. Nitrogen concentration at gauges in Tampa Bay, Charlotte Harbor, and the Indian River Lagoon were reported in terms of concentrations of ammonia (NH_3_), ammonium (NH_4_) and variants of nitrate/nitrite (NO_x_) so observations of DIN, including DIN 1prior, were produced by summing the concentrations of the three components (NH_3_, NH_4_, NO_x_). DIN was not calculated if any of the three components were missing. Monthly river flow rate and precipitation were obtained by averaging daily flow rate and daily precipitation, respectively. Before calculating monthly river flow rate in IR, the flow rates for the north and south prongs of the St. Sebastian River in the Indian River Lagoon were summed to obtain the monthly flow rate of the St. Sebastian River in to the Indian River Lagoon. Additionally, monthly flow rates of the Cedar and Ortega Rivers (oriented similar to the north and south prongs of the St. Sebastian River) which flow in to the north portion of the St. Johns River in northeast Florida (JX) were summed to obtain the total monthly flow rate into the St. Johns River. Monthly flow rates of each river in every estuary were averaged to obtain monthly flow rate of all rivers. Finally, each environmental data type was searched for outliers, and any records outside of the 95^th^ percentile of the distribution were removed.

Observations of DIN, Chlor *a*, river flow, precipitation, Z anomaly, and minimum and maximum T were temporally matched to each sampling event by the month in which each sampling event occurred. DIN was matched to the month prior to the sampling event to account for the time lag between a nutrient loading event and succession of or reduction in phyto- and zooplankton populations (Roelke et al. 1997; Livingston 2001). Observations of at-spawn salinity, at–spawn water temperature, were also temporally matched to each sampling event using the estimated month when the juvenile(s) observed in the sampling event was (were) spawned. Observations of DIN 1prior were matched to the month prior to spawning. Spawning month was estimated by calculating the average age (in months) of juveniles present in each sampling event. The average age was calculated by averaging the total length of juveniles present in each sampling event and applying an age-length relationship for juvenile spotted seatrout (McMichael and Peters 1989). The average age of the fish in each sampling event was then used to estimate an expected spawning month, assuming a pelagic larval duration of two weeks, by subtracting the average age from the sampling month. Therefore, at-spawn salinity, at-spawn water temperature, and DIN 1pior were calculable only when at least one juvenile spotted seatrout was observed in a sampling event.

River flow, precipitation, minimum and maximum T anomalies and the Z anomaly exhibited seasonal lags and were included in each estuary-specific dataset because each variable may influence estuarine circulation and water quality necessary for successful transport, settlement and recruitment of juveniles (Mann and Lazier 1992; Bolle et al. 2009; Dutterer et al. 2013). Average values for each environmental factor during the previous autumn and winter were calculated; the previous autumn encompassed October, November and December of the year prior to each sampling event, and the winter season encompassed January and February of the sampling year. For example, seasonally lagged flow of the closest river was temporally matched to each sampling event by determining the average river flow during October, November, and December of the previous year (thus, previous autumn river flow) and the average river flow during January and February of the current year (thus, winter river flow). The average flow rate of all rivers into each area was also considered on a seasonally lagging timescale. Additionally, the average concentrations of DIN for the two and three months prior (DIN 2prior and DIN 3prior, respectively) to spawning were also considered because of the possible influence of long-term, lower trophic level productivity on spawning stock health, spawning success, and fecundity (Wootton 1998).

Total DIN, Chlor *a*, river flow, at-spawn salinity, at-spawn water temperature, DIN 1prior, DIN 2prior, and DIN 3prior were also spatially matched to each sampling event. Observations of DIN and Chlor *a* from gauges within a 2.5-mile buffer around each sampling event were selected, and the closest observations were matched to the sampling event if the observations were from the same year and month, or month prior for DIN, of the sampling event. Not all sampling events were assigned DIN or Chlor *a* values because many were located more than 2.5 miles away from the closest gauge. The flow rate of the closest river was assigned to each sampling event using the approximate latitudinal and longitudinal coordinates of all river mouths and spatially distinct sampling events in each estuary. Unlike DIN and Chlor *a*, which were matched to the closest sampling event, DIN 1prior, at-spawn salinity and at-spawn water temperature were calculated by averaging the observations of these variables across gauge locations located within probable spawning grounds and during the estimated spawning period of the fish within each sampling event. Observed DIN 2prior and DIN 3prior were assigned to each sampling event only if observations of DIN were available in the vicinity of the spawning grounds and during the two and three months leading up to supposed spawning.

### Data filtering

Structural zeros (true zeros; Zuur et al. 2009) were removed from each estuary-specific dataset to reduce zero inflation using two occurrence rate thresholds. A structural zero was defined as a zero count that occurred at a sampling site because of either timing or habitat reasons. First, a zero count may have occurred because it was too early in the year and new juveniles had yet to settle to the nursery habitat and recruit to the sampling gear. Alternatively, sampling may have been too late in the year so that new juveniles had already undergone ontogenetic habitat shifts and went undetected by sampling. Second, a zero count may have occurred because the structural habitat (for example, bottom vegetation, bottom type, shoreline type) or sampling zone was not and would never be suitable for juvenile spotted seatrout even if it were optimally favorable in terms of other water quality characteristics such as temperature or salinity. A 10% occurrence rate threshold was used for sampling month, and an 85% occurrence rate threshold was used for the structural habitat category. For example, if at least one juvenile spotted seatrout was observed only 10% of the time or less within a certain month across all sampling years within an estuary the sampling events performed during those months were removed. Then, if at least one juvenile spotted seatrout was observed only 15% of the time or less within a specific category of bottom vegetation type, bottom type, shoreline type or sampling zone across all sampling years within an estuary, the sampling events performed in those habitat types were also removed. Finally, sampling events where more than 50 juvenile spotted seatrout were captured were lumped together into a 50+ bin.

### Evaluating environmental predictor importance with XGBoost

The importance of each environmental factor (environmental predictor variable) to the recruitment of YOY spotted seatrout was assessed using gradient boosting regression trees (GBRT) or “gradient tree boosting”, which is a type of classification and regression tree algorithm (CART; Breimen et al. 1993). Classification and regression trees use dominant patterns in predictor variables to partition the response data into homogenous nodes, or leaves, using a series of decisions, i.e., decision rule or decision tree. The resulting decision rule is then used to predict a new series of response data (Breimen et al. 1993; Elith et al. 2008)

Gradient tree boosting is based on principals from classical statistical estimation and machine learning theory (De’ath and Fabricius 2000; Friedman 2001). However, unlike traditional statistical methods that start with a data model, the GBRT algorithm assumes the process by which the response data were produced is complex, unknown, and may have resulted from many inherent variable interactions (Elith et al. 2008). The GBRT algorithm is well-suited for ecological studies exploring predictor importance, particularly those exploring an environment-recruitment relationship, because it can model inherent, multi-level interactions between environmental variables, is generally insensitive to multi-collinearity, can accommodate for outliers and missing observations of predictor variables, and model nonlinearity likely to occur between predictor and response variables following an environmental regime shift (Beamish et al. 1999; De’ath and Fabricius 2000; Polovina 2005). Furthermore, the GBRT algorithm produces a decision rule using *k*-fold cross-validation that can be applied and tested on a withheld portion of data, which is strongly encouraged when testing environment-recruitment relationships (Myers 1998). Gradient tree boosting has been successfully used by fisheries scientists in the past to predict the abundances of spotted seatrout and southern flounder in Texas as well as demersal fish species richness around New Zealand (Leathwick et al. 2006; Froeschke and Froeschke 2011, 2016).

Here, gradient tree boosting was implemented in the R software environment using XGBoost (extreme gradient boosting; R package *xgboost* [Chen et al. 2018]), which is a machine learning method for regression or classification tree boosting that builds many shallow decision trees. Each individual tree describes only a portion of the system but produces a highly accurate rule for prediction when joined in an ensemble (Chen and Guestrin 2016). The XGBoost algorithm is a unique form of gradient tree boosting because decision trees are built to minimize an objective function composed of not only a loss function but also a regularization factor that prevents tree over-fitting (Chen and Guestrin 2016). Additionally, the XGBoost algorithm works in parallel among all available computing cores in combination with the regularization factor to produce a prediction rule faster than most other tree boosting methods (Chen and Guestrin 2016). XGBoost has been used in the ecological literature to forecast bird migration and identify fishing vessel type based on vessel monitoring system trajectories in the East China Sea (Huang et al. 2018; Van Doren and Horton 2018).

Each estuary-specific dataset was randomly shuffled and split into to a training and testing dataset using a 70%/30% split, where 70% of each shuffled dataset was used to train the XGBoost prediction rule, and the remaining 30% was withheld and used to test the decision rule. The 70%/30% ratio is commonly used to split datasets and is supported by validation studies (Guyon 1997; Kuhn et al. 2018). A count Poisson learning objective and a Poisson negative log-likelihood loss objective function were used to create a prediction rule for each estuary-specific dataset. Each XGBoost prediction rule was trained with 10-fold cross-validation to identify the number of trees (*ntree*) that minimized the objective function. Then, using the optimal number of trees and functions within the R *caret* package (Kuhn et al. 2018), the prediction rule was fine-tuned by identifying the optimal combination of hyperparameters that further minimized the objective function for each area. The hyperparameters included the learning rate (*eta*), maximum depth of each tree (*max_depth*), the number of observations in each leaf node of the tree (*min_child_weight*), the minimum loss reduction required to further partition a leaf node on a single tree (*gamma*), the proportion of observed data used by XGBoost to grow each tree (*subsample*), and the proportion of predictor variables used at each level of tree splitting (*colsample_bytree*; DMLC 2016). With the optimal values of the hyperparameters and number of trees, the decision rule was retrained and applied to the withheld testing data to predict a new series of count observations and evaluate the accuracy of the decision rule. The gain value (improvement in accuracy each variable brings to the decision rule, i.e., variable importance) of each environmental predictor variable was also obtained using the XGBoost algorithm.

Next, the combination of environmental predictor variables that produced the lowest mean absolute error (MAE) between the observed and predicted count observations was determined with recursive variable addition. The datasets were stripped of all environmental predictor variables except for salinity and temperature. The data were shuffled, and the training procedure described above was used to determine a decision rule. Additional environmental predictor variables were added to the dataset one at a time, in the order of importance defined by the XGBoost algorithm in the previous step. A decision rule was reproduced after every variable addition and applied to the withheld data to predict a new series of count data. Mean absolute error and mean error (ME) were also calculated after the addition of each variable to explore the accuracy and bias, respectively, of each decision rule (Walther and Moore 2005). Last, the optimal combination of environmental predictor variables for each estuary was used to build a final decision rule; MAE and ME were recalculated, and the predictive performance of each environmental variable was ranked by the XGBoost algorithm.

### Generalized linear models

Generalized linear models (GLMs) were fitted to the same datasets and compared to the XGBoost results. Each dataset was once again shuffled and separated into a testing and training dataset, and the count data were modeled with a Poisson distribution. Step wise model selection was used to identify the environmental predictor variables that explained the most deviance, and only variables that reduced the deviance by 0.5% or more were retained in the final model for each estuary. The final model was applied to the withheld testing data to predict a new series of count data. Lastly, MAE and ME between the predicted and observed data were calculated for each estuary-specific model.

## Results

A total of 902 sampling events in Apalachicola, 2034 sampling events in Cedar Key, 7239 sampling events in Tampa Bay, 6038 sampling events in Charlotte Harbor, 1996 sampling events in northeast Florida, and 6449 sampling events in the Indian River Lagoon along with the associated observations of environmental predictor variables were used to build estuary-specific predictive models of YOY spotted seatrout recruitment (Table 3). Young-of-the-year spotted seatrout were most abundant in Apalachicola, Tampa Bay, Charlotte Harbor, and the Indian River Lagoon where, two YOY spotted seatrout were captured per sampling event on average (Table 3; Figure 2). In contrast, YOY fish were less abundant in Cedar Key and northeast Florida where only one individual was captured per sampling event on average (Table 3; Figure 2).

**TABLE 3.**
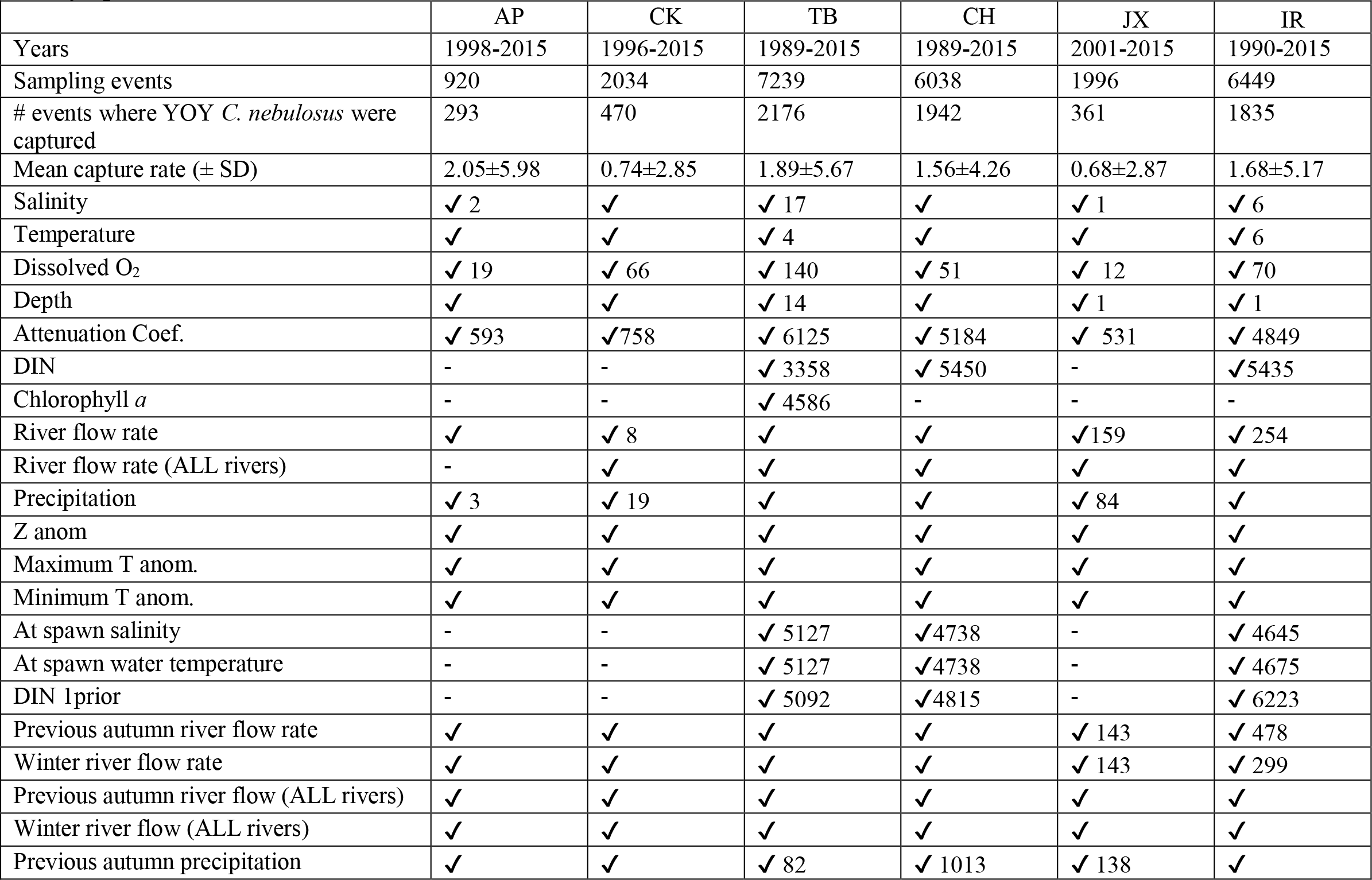
Total number of sampling events within sampling years, number of events where spotted seatrout were captured, mean capture rate, and environmental predictor variables evaluated using the XGBoost algorithm in each estuary: Apalachicola (AP), Cedar Key (CK), Tampa Bay (TB), and Charlotte Harbor (CH), northeast Florida (JX) and the Indian River Lagoon (IR). A check mark indicates whether the variable was used and a number to the right of the check mark is the number of missing observations within the estuary-specific datasets.

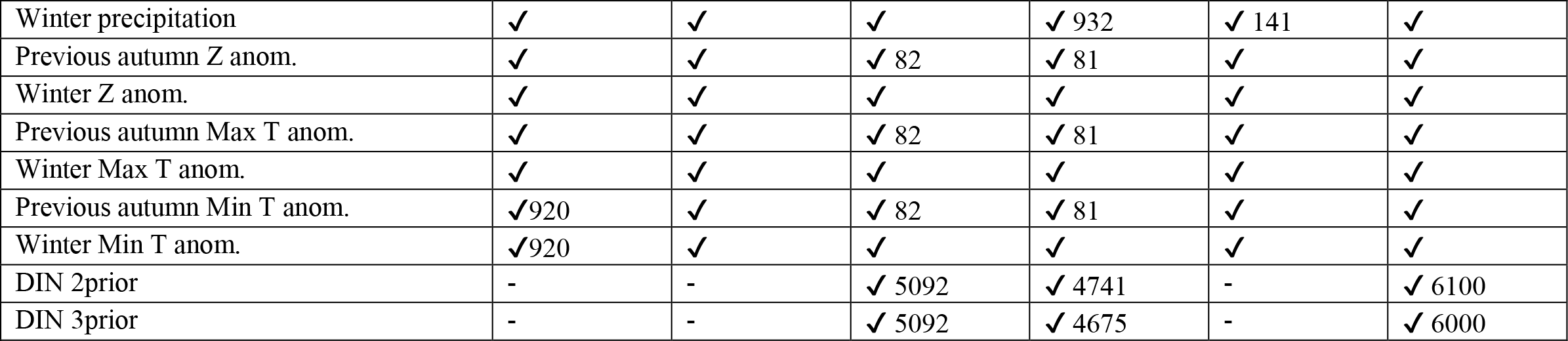

**FIGURE 2.**
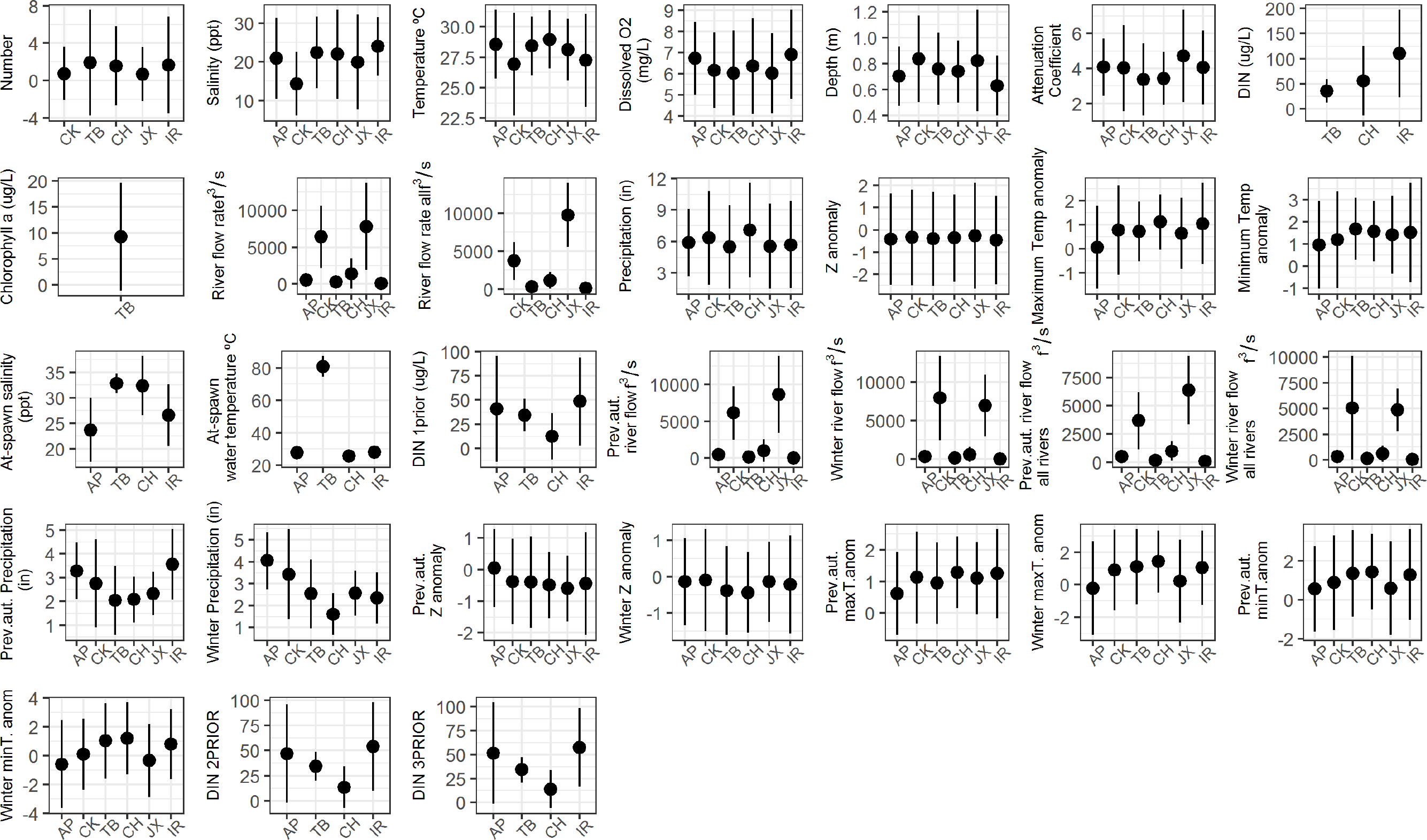
Mean number (± 1 *SD*) of YOY spotted seatrout captured per sampling event, and a summary (mean ± 1 *SD*) of the environmental predictor variables used in the analysis for every estuary: Apalachicola (AP), Cedar Key (CK), Tampa Bay (TB), Charlotte Harbor (CH), northeast Florida (JX), and the Indian River Lagoon (IR).

Observations of salinity, water temperature, dissolved O_2_, depth, precipitation, Z anomaly, and maximum and minimum temperature were available for nearly every sampling event in each estuary (Table 3). River flow rates were missing for a small portion of the sampling events in northeast Florida and the Indian River Lagoon due to either missing river gauge observations or data that did not span the entire sampling timeframe. In contrast, the attenuation coefficient was calculable only 38% of the time, on average, because of frequent sampling in shallow areas where the Secchi disk remained visible when it reached the bottom (Table 3).

Additionally, observations of spatially matched variables including concentrations of DIN and chlorophyll *a*, at-spawn salinity, at-spawn water temperature, and DIN 1prior were infrequent throughout the datasets. Observed DIN was available for only 53% of the sampling events in Tampa Bay and 13% of the sampling events in Charlotte Harbor and Indian River Lagoon on average. Observed chlorophyll *a* was available for only 36% of the sampling events in Tampa Bay. Additionally, at-spawn salinity and at **–**spawn water temperature were matched to only 26% of the sampling events in Tampa Bay, Charlotte Harbor, and the Indian River Lagoon, on average. Additionally, DIN 1prior, DIN 2prior, and DIN 3prior were matched to only 18% of the sampling events in these estuaries, on average.

Salinity, water temperature, dissolved O_2_ and precipitation were generally consistent among areas except for in Cedar Key where the mean salinity and temperature were lowest, t mean depth and the attenuation coefficient varied considerably among all areas (Figure 2). In general, the attenuation coefficient was larger in deeper estuaries as observed in northeast Florida (Figure 2). Additionally, concentration of DIN, river flow rate, at-spawn salinity, at-spawn water temperature, and seasonally lagged precipitation varied widely among areas. The mean concentration of DIN was highest in the Indian River Lagoon and the mean flow rates (including seasonally lagged variants) were highest among rivers in northeast Florida. Both at-spawn salinity and at-spawn water temperature were highest in Tampa Bay and lowest in Apalachicola, while seasonally lagged precipitation was highest in Apalachicola (winter precipitation) and the Indian River Lagoon (previous autumn precipitation) and lowest in Tampa Bay and Charlotte Harbor (previous autumn precipitation and winter precipitation; Figure 2).

The XGBoost algorithm was used to evaluate the predictive performance of 22 environmental variables in Apalachicola, 23 environmental variables in Cedar Key, 29 environmental variables in Charlotte Harbor, 30 environmental variables Tampa Bay, 23 environmental variables in northeast Florida, and 29 environmental predictors in Indian River Lagoon (Table 3). The XGBoost algorithm identified 16 environmental variables that best predicted YOY abundance in Apalachicola, 20 variables for Cedar Key, 28 for Tampa Bay, 26 for Charlotte Harbor, 12 for northeast Florida, and 25 for the Indian River Lagoon (see Table 4 for a list of variables providing gain values greater than 0.01). Salinity, temperature, depth, and dissolved O_2_ exhibited the greatest gains in predictive performance for all areas. River flow during the previous autumn was the most important predictor variable for Cedar Key, and monthly flow of the closest river was among the top ten most important predictor variables for all areas. Precipitation and the Z anomaly were also among the top ten most important predictors for all areas except for northeast Florida and the Indian River Lagoon where the Z anomaly provided a gain of less than 0.05 (Table 4; Figure 3).

**TABLE 4.** Environmental predictor variables that resulted in the lowest mean average error between the observed and predicted counts for each estuary. Predictor variables are listed in order from those resulting in most to least gain as calculated by XGBoost for estuaries on the west coast of Florida; Apalachicola (AP), Cedar Key (CK), Tampa Bay (TB), and Charlotte Harbor (CH)). Note: predictor variables with less than 0.01 gain are not reported here.

| AP |  | CK |  | TB |  | CH |  |
| --- | --- | --- | --- | --- | --- | --- | --- |
| Important Variables | Gain | Important Variables | Gain | Important Variables | Gain | Important Variables | Gain |
| Salinity | 0.17 | Prev. aut. river flow ALL | 0.14 | Salinity | 0.14 | Salinity | 0.19 |
| Temperature | 0.15 | Dissolved O<sub>2</sub> | 0.14 | Dissolved O<sub>2</sub> | 0.11 | Dissolved O<sub>2</sub> | 0.13 |
| Dissolved O<sub>2</sub> | 0.14 | Salinity | 0.11 | Temperature | 0.11 | Temperature | 0.12 |
| Depth | 0.07 | Depth | 0.11 | DIN | 0.08 | Depth | 0.07 |
| Atten. coef. | 0.06 | Temperature | 0.09 | Depth | 0.07 | Precipitation | 0.05 |
| River flow | 0.05 | Year | 0.08 | Chlor *a* | 0.06 | River flow | 0.05 |
| Min. T anom. | 0.05 | Atten. coef. | 0.07 | River flow | 0.05 | River flow ALL | 0.05 |
| Z anom. | 0.04 | River flow | 0.05 | Precipitation | 0.03 | Month | 0.04 |
| Prev. aut. river flow | 0.03 | Precipitation | 0.04 | River flow ALL | 0.03 | Z anom. | 0.04 |
| Precipitation | 0.03 | Z anom. | 0.03 | Z anom. | 0.03 | Prev. aut. river flow | 0.04 |
| Winter Precip. | 0.03 | Max. T anom | 0.03 | Winter river flow | 0.03 | Min. T anom. | 0.03 |
| Max. T anom. | 0.03 | Min. T anom | 0.02 | Max. T anom. | 0.03 | Winter river flow | 0.03 |
| Winter river flow | 0.02 | River flow ALL | 0.02 | Winter river flow | 0.03 | DIN | 0.02 |
| Prev. aut. Precip. | 0.02 | Month | 0.02 | Atten. coef. | 0.02 | Max T. anom. | 0.02 |
| Year | 0.01 | Winter river flow | 0.01 | Month | 0.02 | Atten. coef. | 0.02 |
| Month | 0.01 | Winter max T anom. | 0.01 | Prev. aut. river flow | 0.02 | Winter max T. anom. | 0.01 |
|  |  | Prev. aut. max T anom | 0.01 | Min. T anom. | 0.01 | Prev. aut. Z anom. | 0.01 |
|  |  | Prev. aut. Z anom | 0.01 | Prev aut. Z anom. | 0.01 | Year | 0.01 |
|  |  |  |  | Year | 0.01 |  |  |
|  |  |  |  | Winter river flow | 0.01 |  |  |

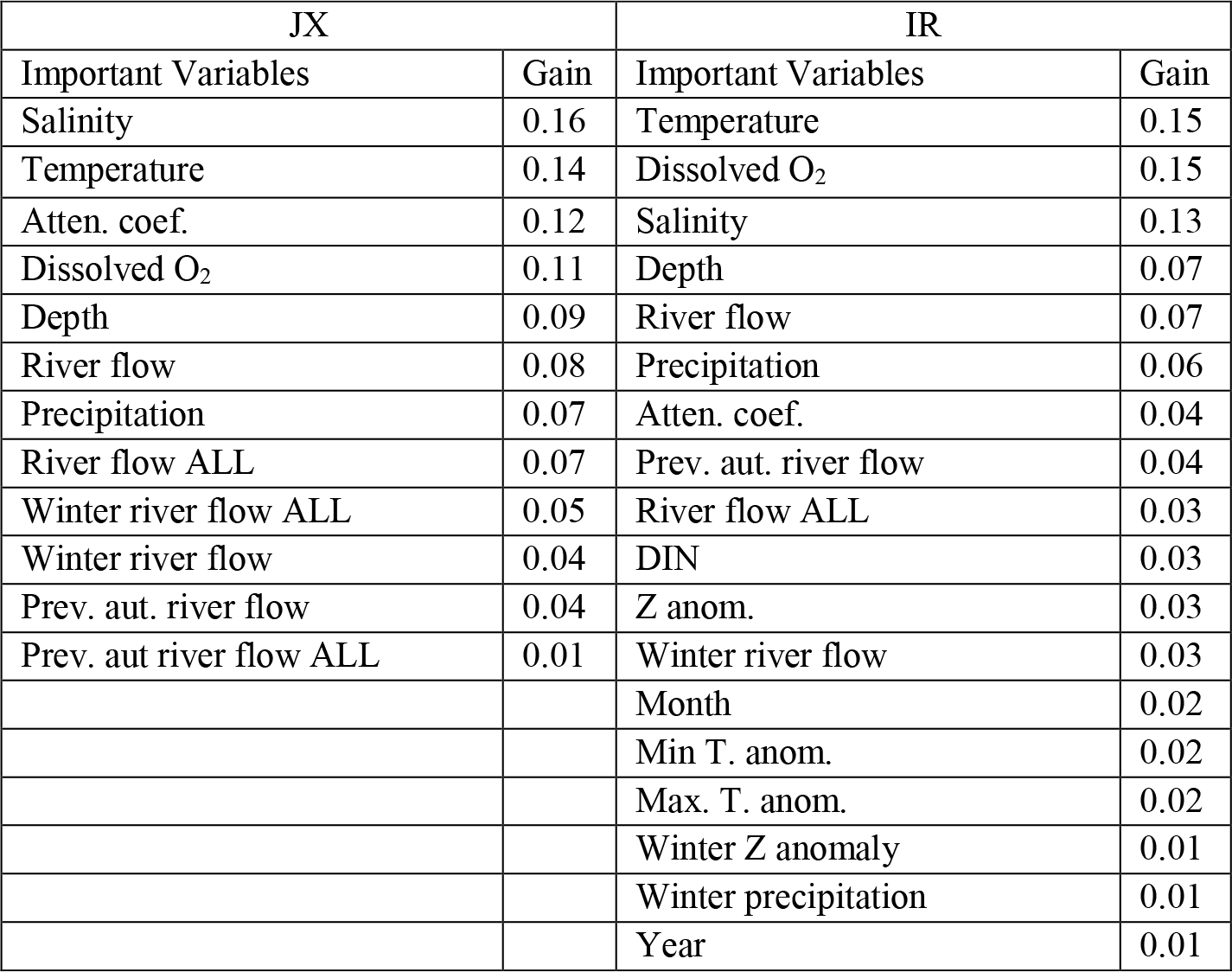

**FIGURE 3.**
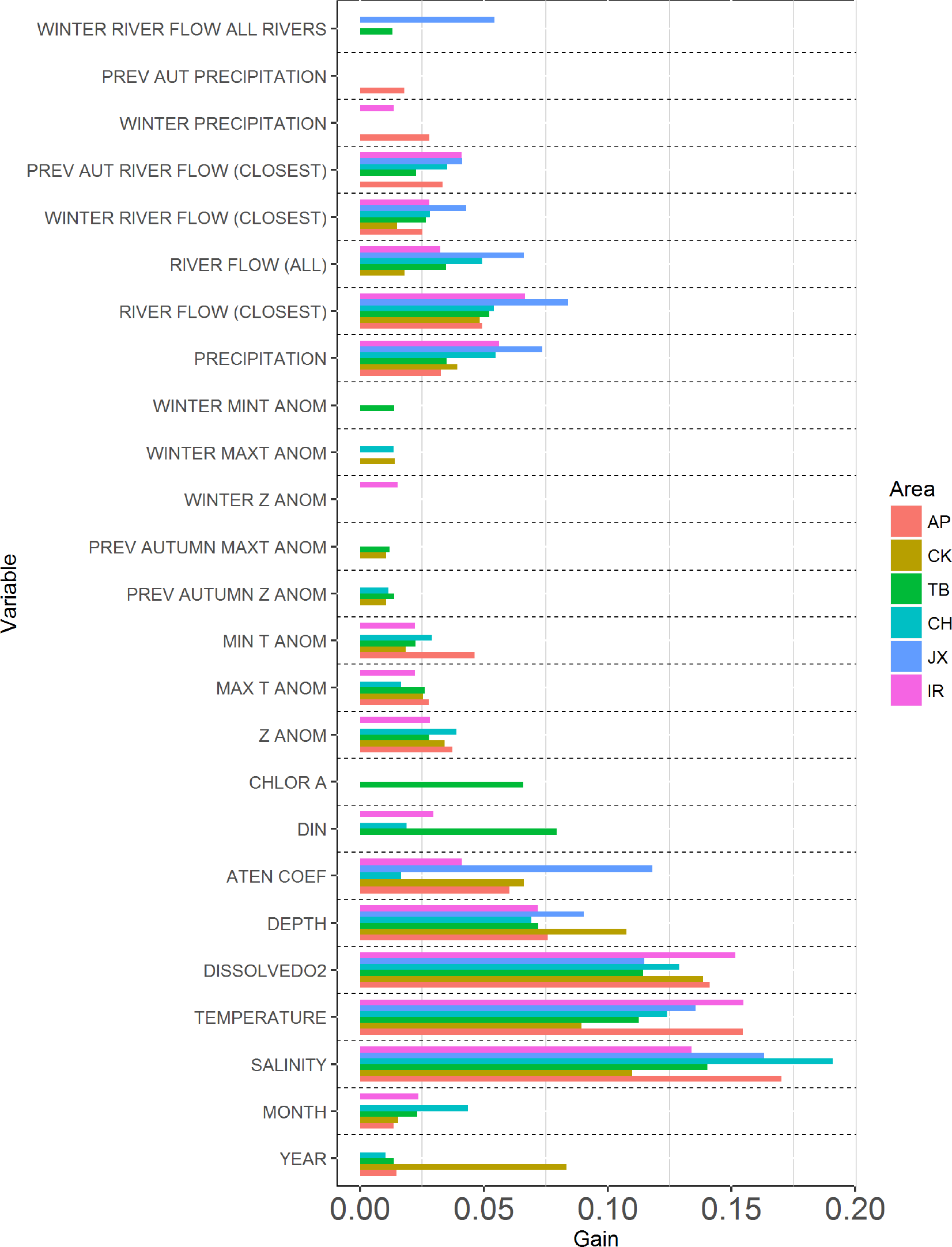
Importance of environmental predictor variables by estuary: Apalachicola (AP), Cedar Key (CK), Tampa Bay (TB), Charlotte Harbor (CH), northeast Florida (JX), and the Indian River Lagoon (IR).

In Apalachicola, the top ten most important environmental predictors were salinity (gain = 0.17), water temperature (gain = 0.15), dissolved O_2_ (gain = 0.14), depth (gain = 0.07), attenuation coefficient (gain = 0.06), river flow (gain = 0.05), minimum temperature anomaly (gain = 0.05), Z anomaly (gain = 0.04), average river flow during the previous autumn (gain = 0.03), and precipitation (gain = 0.03). In addition, precipitation during the previous autumn and winter seasons, maximum temperature anomaly, winter river flow, and sampling year and month provided a combined gain of 0.12 (Table 4; Figure 3). In Cedar Key, the top ten most important environmental predictors were average river flow during the previous autumn (gain = 0.14), dissolved O_2_ (gain = 0.14), salinity (gain = 0.11), depth (gain = 0.11), temperature (gain = 0.09), year (gain = 0.08), attenuation coefficient (gain = 0.07), river flow (gain = 0.05), precipitation (gain = 0.04), and Z anomaly (gain = 0.03). In addition, maximum and minimum temperature anomalies, river flow of all rivers, sampling month, average river flow during the previous autumn and winter, average maximum temperature anomaly during the previous autumn and winter, and average Z anomaly during the winter provided a combined gain of 0.13 (Table 4; Figure 3). In Tampa Bay, the top ten most important environmental predictors were salinity (gain = 0.14), dissolved O_2_ (gain = 0.12), water temperature (gain = 0.11), concentration of DIN (gain = 0.08), depth (gain = 0.07), concentration of chlorophyll *a* (gain = 0.06), river flow (gain = 0.05), precipitation (gain = 0.03), river flow of all rivers (gain = 0.03), and the Z anomaly (gain = 0.03). In addition, minimum and maximum temperature anomalies, average river flow during the previous autumn and winter, the attenuation coefficient, sampling year and month, average minimum temperature anomaly during winter, average Z anomaly and maximum temperature anomaly during the previous autumn provided a combined gain of 0.19 (Table 4; Figure 3). In Charlotte Harbor, the top ten most important environmental predictors were salinity (gain = 0.19), dissolved O_2_ (gain = 0.13), temperature (gain = 0.13), depth (gain = 0.07), precipitation (gain = 0.05), river flow (gain = 0.05), river flow of all rivers (gain = 0.05), sampling month (gain = 0.04), Z anomaly (gain = 0.04), and average river flow during the previous autumn (gain = 0.04). In addition, minimum and maximum temperature anomalies, average river flow during winter, concentration of DIN, the attenuation coefficient, average maximum temperature anomaly during winter, average Z anomaly during the previous autumn and sampling year provided a combined gain of 0.15 (Table 4; Figure 3).

In northeast Florida, the top ten most important environmental predictors were salinity (gain = 0.16), temperature (gain = 0.14), attenuation coefficient (gain = 0.12), dissolved O_2_ (gain = 0.11), depth (gain = 0.09), river flow (gain = 0.08), precipitation (gain = 0.07), river flow of all rivers (gain = 0.07), and average river flow during winter (all and closest, gain = 0.05 and gain =0.07, respectively). In addition, average river flow of the closest and all rivers during the previous autumn provided a combined gain of 0.05 (Table 4; Figure 3). In the Indian River Lagoon, the top ten most important environmental predictors were temperature (gain = 0.15), dissolved O_2_ (gain = 0.15), salinity (gain = 0.13), depth (gain = 0.07), river flow (gain = 0.07), precipitation (gain = 0.06), the attenuation coefficient (gain = 0.04), average river flow during the previous autumn (gain = 0.04), river flow of all rivers (gain = 0.03), and concentration of DIN (gain = 0.03). In addition, the Z anomaly, average river flow, precipitation, and maximum temperature anomaly, Z anomaly during winter, maximum temperature anomaly, minimum temperature anomaly, sampling year and month provided a combined gain of 0.15 (Table 4; Figure 3).

The mean absolute error between the observed and predicted count data in all estuaries indicated good predictive performance by the XGBoost decision rule. The XGBoost algorithm produced a decision rule predicted the count data distribution more accurately that the GLMs (MAE = 2.15 vs 3.19 for AP; 0.90 vs 1.08 for CK; 2.75 vs 3.23 for TB; 1.56 vs. 1.99 for CH; 0.53 vs 0.71 for JX; 1.96 vs 2.37 for IR; Table 5). However, the distributions were slightly more biased (higher ME) than those produced by the GLMs (0.90 vs −0.94 for AP; 0.53 vs 0.09 for CK; 1.06 vs 0.04 for TB; 0.60 vs −0.18 for CH; 0.48 vs −0.02 for JX; 0.8 vs −0.11 for IR; Table 5).

**TABLE 5.** Sample sizes of the training and testing datasets, mean absolute error (MAE), and mean error (ME) between observed and predicted counts produced by the XGBoost algorithm (XGB) and generalized linear model (GLM). Estuaries are Apalachicola (AP), Cedar Key (CK), Tampa Bay (TB), Charlotte Harbor (CH), Indian River Lagoon (IR), and northeast Florida (JX).

| Estuary | Model | Sample size<br>(N)<br>train; test | MAE | ME |
| --- | --- | --- | --- | --- |
| AP | XGB | 644; 276 | 2.45 | 0.90 |
|  | GLM | 554; 237 | 3.19 | -0.94 |
| CK | XGB | 1424; 610 | 0.90 | 0.53 |
|  | GLM | 580; 248 | 1.08 | 0.09 |
| TB | XGB | 5067; 2172 | 2.38 | 1.06 |
|  | GLM | 1714; 735 | 3.23 | 0.04 |
| CH | XGB | 4227; 1811 | 1.84 | 0.60 |
|  | GLM | 3023; 1296 | 1.99 | -0.18 |
| JX | XGB | 1397; 599 | 0.79 | 0.48 |
|  | GLM | 1090; 467 | 0.71 | -0.02 |
| IR | XGB | 4514; 1935 | 2.07 | 0.8 |
|  | GLM | 4127; 1769 | 2.37 | -0.11 |

## Discussion

This study synthesized the results of a large body of research on the environmental factors affecting spotted seatrout and ranked the importance of environmental predictor variables using a new gradient tree boosting algorithm. The relative importance of each environmental predictor variable was consistent among areas, and any differences are reflective of estuary-specific differences in geomorphology, circulation patterns and habitat characteristics. Salinity, temperature, dissolved O_2_, and depth were all among the five most important predictor variables for all estuaries. In addition, river flow was among the seven most important environmental predictor variables in all estuaries, except for in Cedar Key where it was the most important variable. Precipitation and drought were among the ten most important predictor variables in all estuaries.

Spotted seatrout are euryhaline species and have been found in waters ranging widely in salinity and temperature (Bortone 2003). However, rapidly changing water conditions can impart lethal osmoregulatory stress on larvae and juveniles as well as limit metabolic activity and oxygen transfer ability and, therefore, strongly influence the distribution and abundance of juveniles and adults (Banks et al. 1991; Alsuth and Gilmore 1994; Holt and Holt 2003; Whitfield and Harrison 2003; Wuenschel et al. 2004). Salinity and temperature conditions also greatly influence spawning success by affecting fertilization and hatching rates, egg buoyancy, and the metabolic rates of reproductive adults (Holliday 1969; Fry 1971; Gray et al. 1991; Holt and Holt 2003; Kupschus 2004). In addition, the concentration of dissolved oxygen affects growth, survival, and respiration of larval and juvenile fishes and can influence predator-prey interactions by altering the vertical distribution of species (Breitburg et al. 1994, 1999; Secor, D.H.; Gunderson 1998). Our results echo those of Froeschke and Froeschke (2011), who also used gradient boosting to explore a suite of environmental predictor variables for spotted seatrout populations in Texas and found that salinity, temperature, turbidity, and dissolved oxygen were the most important variables among nine estuary populations.

The influx of freshwater from rivers and other point sources can significantly alter the salinity, temperature, dissolved oxygen, turbidity, light availability, and nutrient conditions within estuary systems thereby affecting the abundance and distribution of juvenile fishes (Kimmerer 2002; Halliday et al. 2008; Gillanders et al. 2012; Whaley et al. 2016; de Mutsert et al. 2017). Variability in freshwater flow can also impact spawning success and the subsequent shoreward larval transport and settlement to essential nursery habitat by altering the salinity gradient, stability of the halocline, and estuary circulation (Drinkwater and Frank 1994; Pineda et al. 2007; Gillson 2011). In addition, freshwater flow can impact the quality and distribution of nursery habitat as salinity intrusions or even overall habitat loss can occur during reduced flow or drought conditions (Drinkwater and Frank 1994; Gillson 2011; Whaley et al. 2016). Our results support the findings of previous research on spotted seatrout and freshwater flow in Florida. Purtlebaugh and Allen (2010) found a positive relationship between freshwater flow from the Suwannee River and abundance of YOY spotted seatrout. In addition, Matheson et al. (2003) and Flaherty et al. (2015) found a positive relationship between YOY spotted seatrout abundance and freshwater flow in Tampa Bay.

Flow rate of the closest river offered greater gain than the average flow rate of all rivers for each estuary with one exception. Average nutrient and water quality conditions in estuaries are influenced by water circulation which is controlled by tides, winds, and the salinity gradient between the less saline freshwater and the resident salt water (Mann and Lazier 2013). As such, estuarine water quality may remain heterogeneous even when the average flow rate of all rivers increases. For example, the Indian River Lagoon is long and narrow and has relatively sluggish wind driven circulation with limited tidal flushing. Therefore, while portions of the Indian River Lagoon may suddenly receive increased nutrient-laden freshwater, other areas and the surrounding flora and fauna may remain unaffected (Sigua and Tweedale 2003). Additionally, major salinity gradients limit the circulation among Tampa Bay’s four major segments, thus, the nutrient-laden freshwater flowing into the northern portion of the bay may not necessarily result in a net increase in nutrients in the lower portions of the bay (Greening and Janicki 2006). Therefore, the flow rate of the river closest to the capture location or home range of a pre-recruit or juvenile spotted seatrout is more influential to the surrounding water quality and nutrient conditions than the average flow rate of all rivers to an estuary.

In contrast, the average flow of both rivers in Cedar Key (Suwannee River and Waccasassa River) during the previous autumn was the most important environmental predictor variable. We hypothesize that this result occurred because the average flow rate of the Suwannee River (which flows at the second highest rate of all rivers in Florida) from 1996 through 2015 (8175 f^3^/s) was nearly 33 times that of the Waccasassa River (245 f^3^/s) therefore, the juvenile spotted seatrout that were captured closer to the mouth of the Waccasassa River in the southern portion of the Cedar Key sampling area were influenced by freshwater from both the Suwannee River and the Waccasassa River. The results from Purtlebaugh and Allen (2010) corroborate our findings. They found that spotted seatrout in the southern portion of the Cedar Key sampling area were strongly influenced by freshwater from the Suwannee River. However, the seasonal lag contrasts with that identified by Purtlebaugh and Allen (2010), who found that river flow from March through May explained most of the variation in spotted seatrout abundance. This current study evaluated average flow rates from only October through December and January through February. Our results may be more comparable had we evaluated average flow rates from March through May, as evaluated in Purtlebaugh and Allen (2010). Average flow rates of the Suwannee and Waccasassa Rivers peak between February and April, and a smaller peak occurs between August and October (Purtlebaugh and Allen 2010); thus, it is unclear why the average winter flow rate, which includes the month of February in this study, was less important. One hypothesis is that the influx of nutrient-laden freshwater in autumn sets the stage for the productivity of the Cedar Key estuary in the coming year by affecting upper level productivity and spawning stock health. However, future studies are necessary to explore a link between autumn conditions and juvenile spotted seatrout abundance in Cedar Key.

In general, river flow is strongly influenced by precipitation and drought. Although decreased precipitation during drought conditions may result in decreased river flow rates, the two variables may not always correlate. In Florida, minimum flow and water levels of major water resources are maintained by water management districts to mitigate the effects of water withdrawals, and conversely, to prevent flooding (Olexa et al. 2017). While patterns of increased precipitation or short-term drought conditions may alter salinity gradients or the distribution and availability of essential nursery habitat, precipitation and drought cycles are not directly responsible for the influx of nutrients to the estuaries. Therefore, both precipitation and drought, while among the ten most important variables, were less important than river flow in all estuaries.

The importance of several environmental predictors varied among estuaries. In Tampa Bay, total dissolved inorganic nitrogen as well as chlorophyll *a* were more important than river flow, as excess nutrients from agricultural runoff or increased river flow stimulate plant growth and can induce oxygen-depleting, algal blooms that may result in hypoxic or anoxic conditions. In addition, increased nutrients can induce a shift from macrophyte to phytoplankton or algal-based communities (Sigua and Tweedale 2003; Greening and Janicki 2006). For example, increased nutrient loading in Tampa Bay during the late 1970’s and early 1980’s was responsible for the precipitous decline in seagrass, which is important habitat for juvenile spotted seatrout (Flaherty-Walia et al. 2015). While highly important to juvenile spotted seatrout abundance in Tampa Bay, DIN was of low importance in Charlotte Harbor and the Indian River Lagoon. This discrepancy is possibly a result of disproportionate number of observations among estuaries. While more than half (54%) of the sampling events in Tampa Bay had associated DIN observations, only 14% of the sampling events in the Indian River Lagoon and 10% of the sampling events in Charlotte Harbor had such observations. While the XGBoost algorithm can infer missing values to a certain degree (Chen and Guestrin 2016), the paucity of observations in Charlotte Harbor and the Indian River Lagoon may have exceeded the ability of the XGBoost algorithm to identify a decision rule based on limited observations. The predictive performance of the XGBoost decision rule tuned to varying levels of missing data requires further investigation.

Light attenuation, which is partially a function of phytoplankton biomass, turbidity and eutrophication affects the distribution and abundance of seagrass, which is important habitat structure for pre-recruits and juvenile fish (Flaherty-Walia et al. 2015). In addition, light availability affects the distribution and abundance of benthic macro-algae which is a basal resource for many estuarine species including juvenile spotted seatrout (Burghart et al. 2013). An attenuation coefficient was not calculated if the Secchi disk was still visible at the bottom of the water column; thus, sampling events in deeper, less clear water were more likely to have an associated attenuation coefficient than those in shallower, clearer water. For example, the attenuation coefficient was the third most important variable in northeast Florida where it was missing for only 26% of the sampling events. In contrast, in shallower or clearer estuaries such as Tampa Bay and Charlotte Harbor, only an average of 15.5% of the sampling events had associated attenuation coefficients, thus, the XGBoost algorithm had few observations to inform the decision rule training process. Direct measures of the factors contributing to light attenuation in estuaries including turbidity, phytoplankton biomass or water color, should be evaluated in future predictive studies to provide a more complete series of environmental predictors variables and more data with which to tune a decision rule.

Among all estuaries, maximum and minimum air temperature anomalies and their seasonally lagged versions were of little importance. The temperatures of water and air change at different rates, so seasonally anomalous air temperature conditions are not immediately reflected by the water temperature. Air temperature anomalies have little predictive power when evaluated on a monthly and or seasonal timeframe but may have greater predictive power if they are considered on a wider timeframe. This hypothesis requires further investigation. In addition, temperature, salinity, and DIN at estimated time of spawning were not included in the final XGBoost decision rule for any estuary dataset on which they were evaluated. While temperature and salinity affect spawning success in spotted seatrout (Kupschus 2004), the relationship between these variables and juvenile abundance was not captured by the XGBoost decision rule, which is likely a result of the limited observations and generalizations in supposed spawning areas, timing and average environmental conditions in spawning grounds.

While the tolerance of the XGBoost algorithm to missing values requires further investigation, the XGBoost algorithm produced a decision rule that predicted the count distribution more accurately than a traditional GLM. The use of gradient boosting in the fisheries and marine ecology literature has increased over the past ten years. A Google Scholar search to find articles with the words “fish”, “marine” and the exact phrase “gradient boosting” published between 2008 and 2012 yielded 142 results, while the same search for articles published between 2013 and 2018 yielded 316 results. As the XGBoost algorithm is new to the machine learning literature, there are few published studies in the fields of fisheries or marine ecology that detail XGBoost usage. A Google Scholar search to find articles with the words “fish”, “marine”, and “XGBoost” published between 2014 and 2018 yielded 6 results, but the results of this study show that this algorithm has strong predictive power when evaluating species abundance and identifying important environmental factors. It is strongly encouraged that future research explore the applicability of the XGBoost algorithm to other topics in marine and fisheries science and compare its performance to that of other statistical methods.

This study showed that the abundance of pre-recruit, juvenile spotted seatrout in spatially distinct estuaries is influenced by nearly the same set of environmental predictors and each predictor was similarly important. Water characteristics including salinity, temperature, concentration of dissolved oxygen, and water quality are most important. River flow is also highly important. While the goal of this study was to evaluate and rank the importance of every possible environmental variable, toxic algal blooms and seagrass acreage can also affect the growth and mortality of juveniles (Flaherty and Landsberg 2011; Flaherty-Walia et al. 2015). These variables were not included in this study, and future research should evaluate their importance in a decision rule framework. The results of this study can inform future assessment and management efforts for spotted seatrout. The environment can strongly affect population productivity, so the inclusion of highly important environmental variables in future stock assessments should be evaluated. Such inclusion may explain a portion of the process error common to many modern stock assessments and afford more accurate estimates of important population parameters and management reference points for this species.

## Acknowledgments

The survey data for spotted seatrout were provided courtesy of the Fisheries Independent Monitoring program of the Florida Fish and Wildlife Conservation Commission’s Fish and Wildlife Research Institute (FWRI) in St. Petersburg, FL. The author would like to acknowledge Dr. Robert Muller and Shanae Allen of FWRI, who provided suggestions on methodology that greatly helped this manuscript. Special thanks go to Ms. Sue Connors and Dr. Margaret Guyette of the St. Johns River Water Management District who assisted with data procurement for the Indian River Lagoon. This manuscript was completed while E.H.S. was employed by FWRI.

